# A comparative analysis of network mutation burdens across 21 tumor types augments discovery from cancer genomes

**DOI:** 10.1101/025445

**Authors:** Heiko Horn, Michael S. Lawrence, Jessica Xin Hu, Elizabeth Worstell, Nina Ilic, Yashaswi Shrestha, Eejung Kim, Atanas Kamburov, Alireza Kashani, William C. Hahn, Jesse S. Boehm, Gad Getz, Kasper Lage

## Abstract

Heterogeneity across cancer makes it difficult to find driver genes with intermediate (2-20%) and low frequency (*<*2%) mutations^1^, and we are potentially missing entire classes of networks (or pathways) of biological and therapeutic value. Here, we quantify the extent to which cancer genes across 21 tumor types have an increased burden of mutations in their immediate gene network derived from functional genomics data. We formalize a classifier that accurately calculates the significance level of a gene’s network mutation burden (NMB) and show it can accurately predict known cancer genes and recently proposed driver genes in the majority of tested tumours. Our approach predicts 62 putative cancer genes, including 35 with clear connection to cancer and 27 genes, which point to new cancer biology. NMB identifies proportionally more (4x) low-frequency mutated genes as putative cancer genes than gene-based tests, and provides molecular clues in patients without established driver mutations. Our quantitative and comparative analysis of pan-cancer networks across 21 tumour types gives new insights into the biological and genetic architecture of cancers and enables additional discovery from existing cancer genomes. The framework we present here should become increasingly useful with more sequencing data in the future.

## Introduction

Understanding which genes have causative roles in cancer will lay the basis for precision medicine, diagnostics and the development of therapeutics. The recent revolution of cancer-sequencing studies has identified many new classes of cancer genes, hereby pointing to new tumour biology^2,3^. From these studies it is also clear that high-frequency cancer genes have already been identified to a large extent, but that we are still missing many of the genes hidden in the long tail of genes mutated at low frequencies^1^ of cancer genomes that can help us to understand the genetic requirements of tumour formation. These genes may also point to new pathways or networks of therapeutic or diagnostic relevance.

Because genes collaborate in functional molecular networks^2,4,5^, network-based analyses have been applied widely in cancers to provide a molecular stratification of cancer patients^4^, to predict disease outcome^6,7^, to understand tumourigenesis^8^ and tumour-inducing viruses^9^, to predict carcinogenicity of chemical compounds^10^, and to prioritize damaging effects of cancer mutations^11^. Several approaches have also been used to identify oncogenic pathway modules^12,13^.

Here, we sought to make four alternative analyses: *First*, to make a comprehensive quantification and comparison of the extent to which the functional gene neighborhood of a large set of established cancer genes, as well as genes emerging from the recent onslaught of cancer sequencing studies across many different tumour types, have an increased network mutation burden (NMB) using pan-cancer mutation information from Lawrence et al. (Ref. 1). Gene-based tests like MutSigCV^1^ calculate significances by aggregating all mutations observed in a cohort across a gene and tools like MutSigCL^1^ and MutSigFN^1^ look at clustering of mutations and enrichment at likely functional sites, respectively (we will refer to these three tools as the ‘MutSig suite’ hereafter). In contrast, we wanted to provide an accurate statistic for the burden of mutations across an index genes’ entire functional neighborhood (defined here as all genes that have a direct connection to the index gene in the high confidence protein-protein interaction network InWeb^14^, while excluding the gene itself), and to use this information to make statistically robust classifications of known and recently proposed cancer genes. *Second*, we wanted to complement and expand current sequencing results by testing whether the NMB can provide an alternative approach to predict new putative cancer genes. *Third*, to ask whether the identification of a putative cancer gene by proxy of its NMB could potentially overcome some of the challenges of finding likely cancer genes with intermediate and low frequency mutations in tumours. *Fourth*, to test the ability of our method to suggest molecular clues as to what drives cancer in patients with no known driver mutations.

Our underlying hypothesis is that summing up the mutations in the neighborhood of a gene can increase the signal to find important cancer genes that are mutated too rarely to be discovered by current gene-based approaches such as the MutSig suite. Towards this aim we develop an accurate statistical method to calculate the significance level of the mutation burden of the functional gene network of cancer genes. We use this method to comprehensively explore and compare the genetic and functional architecture of cancer networks in 21 tumour types. For 60% of the cancer types we test, the NMB approach can accurately recapitulate established driver genes based on either the mutations observed in the corresponding tumour-type or by exploiting pan-cancer mutation rates and significances. Using NMB, we predict 62 putative cancer genes. The NMB approach finds proportionally more (4x) low-frequency putative cancer genes than gene-based tests and suggests molecular clues in patients with, for example, colorectal cancer, kidney clear cell carcinoma and prostate cancer that do not have mutations in established driver genes.

Overall we harness the fundamental wiring of genes into functional networks to develope a robust statistical framework that complements gene-based tests to generate new hypotheses about driver-gene candidates. Our approach should become increasingly useful as more cancer genomes are sequenced in the future.

## Results

### Network Mutation Burden distinguishes true cancer genes

For a given index gene the NMB is formalized into a score that reflects the empirical probability of the observed mutation signal aggregated across its first order biological network, excluding the index gene itself, while normalizing for the number of genes in this network (**Methods**). For this analysis we used the high confidence subset of InWeb^14^ (a quality controlled protein-protein interaction network) as it has already been used in dozens of genetic studies including the 1000 Genomes Project^15^). We confirmed that the NMB accurately calculates the significance level of the mutation burden in the neighborhood of an index gene, based on the fact that the majority of genes fit the null hypothesis and lie on the diagonal in a Q-Q plot (**Supplementary Figure 1**). By testing several permutation methods and alternative ways of calculating the NMB we obtain similar results, supporting the bi-ological and statistical robustness of the NMB concept and of our chosen approach (**Supplementary Note 1** and **Supplementary Figure 2**).

To test if the NMB score can accurately classify cancer genes, we curated four sets of genes linked to cancer and randomly chose a set of genes for control purposes (**Supplementary Note 2** and **Supplementary Table 1**). Briefly, the sets were named Tier 1-5, where Tier 1 genes are well established, or “classic”, cancer genes from the Catalogue of Somatic Mutations in Cancer, or Cosmic, (e.g. *TP53*, *BRCA1*, and *BRAF*); Tier 2 genes are a set of more recently identified cancer genes from the Sanger Gene Census dataset in some cases with functional support (e.g. *MLL2*, *CDK12*, and *GATA2*); Tier 3 genes are recently emerging cancer genes that have been identified using conservative statistics in cancer sequencing studies, but where the biological connection to known cancer pathways is often unclear (e.g. *ING1*); Tier 4 are suspected cancer genes with solid, but in some cases not entirely conclusive statistical evidence from cancer sequencing studies (e.g. *EIF2S2*), and Tier 5 is a random set of genes not linked to cancers included as a control for cryptic confounders in our analysis.

We tested whether the NMB score could accurately distinguish Tiers 1-5 from all genes covered by interactions in the InWeb database that are not in Tiers 1-5 (which is conservative as many of the genes not in Tier 1-5 may be genuine cancer genes that have not yet been discovered). We were able to distinguish genes in Tiers 1, 2 and 3 from other genes in InWeb with an area under the receiver operating characteristics curve (AUC) of 0.86, 0.67, 0.75, respectively (**Fig. 1a**, Adj. P *<*0.05 for each of these AUCs, using permuted networks **Supplementary Figure 2**). The AUC for Tier 4 is insignificant (nominal P = 0.13), consistent with the lower confidence that Tier 4 represents true cancer genes. Tier 5 was indistinguishable from other InWeb genes (**Fig. 1a**, AUC 0.49, nominal P = 0.81) as expected of a random set of genes. The same analysis was repeated for all known pan-cancer genes (**Supplementary Table 2**), yielding an AUC of (0.70, Adj. P *<*0.05, not shown), where the lower AUC is to be expected as it is a mix of genes from Tiers 1-4, with proportionally more genes from Tier 4 (**Supplementary Note 3**).

Tier 1 genes are generally very well studied and have been known for many years to be key drivers of tumourigenesis through their participation in cellular networks involved in DNA repair, cell cycle, proliferation and apoptosis. They also distinguish themselves by having the highest mutation frequencies and most significant mutations across patients in 21 tumour types (**Supplementary Figure 3**). Although we do not observe any correlation between the number of interactions (termed “degree” hereafter) of a gene, its NMB significance, or its Tier membership (**Supplementary Figure 4**) we quantified the potential effect of study bias in Tier 1 genes on our results by canceling their influence on the NMB calculation and repeating the analysis (**Fig. 1b** and discussed in detail in **Supplementary Note 4**). Although the ability to classify Tier 1 genes is slightly decreased, there was no general effect on the accuracy of the NMB classification on Tiers 2, 3 and 4, reflecting that the signal for genes in Tiers 2, 3 and 4 is independent of the Tier 1 genes and not affected by study bias or “knowledge contamination”. This observation is corroborated by the fact that genes in Tiers 2, 3, and 4 have significant overall connectivity to other genes in Tiers 2-4 and not just to genes in Tier 1, hence the NMB signal in these tiers is independent of Tier 1 genes (**Supplementary Figure 5**).

**Figure 1.**
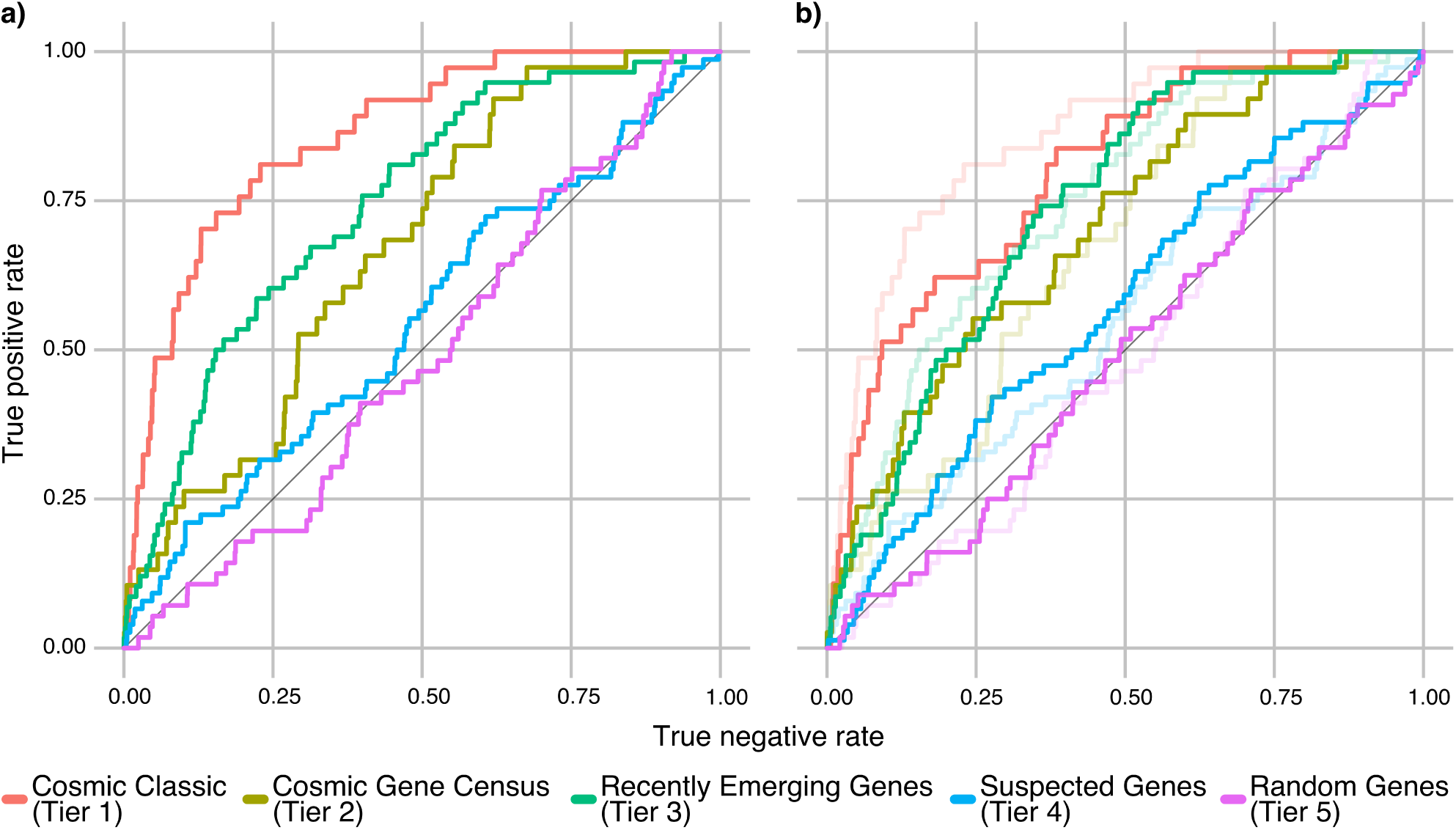
Network mutation burden distinguishes true cancer genes and is unaffected by potential “knowledge contamination”. **a)** Genes from Tiers 1-5 are in this analysis defined as true positives and all other genes in InWeb (12,500 not in Tiers 1-5) as true negatives. Genes in Tier 5 are random genes that are included as a control under the assumption that this gene set should fit the null hypothesis and have an AUC 0.5. NMB scores for all genes are calculated using pan-cancer significances (described in **Methods**), and used as a classifier. The areas under the receiver operating characteristics curve (AUCs) for Tiers 1,2,3, and 4 are 0.87, 0.67, 0.75, 0.55 (Adj. P *<*0.05 for Tiers 1,2 and 3). Genes from Tier 5 fit the null hypothesis (AUC 0.49, nominal P = 0.75). **b)** Removing the effect of Tier 1 genes on the analysis has some impact on the classification of Tier 1 genes themselves, but not on the other Tiers (fully drawn lines represent the results excluding Tier 1 genes shaded lines the original results for comparison). This illustrates that well-established cancer genes are not driving the ability of the NMB score to correctly classify cancer genes.

### Predicting sixty-two NMB-imputed cancer genes

We declared genes with a pan-cancer NMB significance at a false discovery rate (FDR q *<*= 0.1) to be NMB-imputed cancer genes (**Fig. 2** and **Supplementary Table 3**). In addition to the pan-cancer results, we performed the NMB analysis on mutation data from each of the 21 tumours independently and also declared genes with q *<*= 0.1 in each of the individual cancers NMB-imputed cancer genes (**Fig. 2**). We use the terminology ’NMB-imputed cancer genes’ to emphasize the fact that the signal in our analysis is based on the network neighborhood of the gene and not coming from the gene itself. Consequently, the NMB significances are independent, and fully complementary, to the gene-based methods (such as the MutSig suite).

We further constructed a dataset of all unique genes that were significant in the pan-cancer analysis pooled with those significant in at least one of the 21 tumour types and called this set the NMB5000 set. In contrast to earlier observations of mutation significances1, we do not detect more genes when using the tumour-specific approach, which highlights the benefit of aggregating patients with different tumour types into one pan-cancer cohort to increase the statistical power to unravel new cancer networks and biology. We note that although we are pooling data from 22 tests, the vast majority of significant genes (85%) come from the pan-cancer analysis alone so the true false discovery rate of this pooled set should only be marginally higher than 0.1.

Our NMB5000 set comprises 62 NMB-imputed cancer genes. To assess their validity and to link these genes to known cancer biology we carried out a comprehensive literature review (see **Supplementary Note 5** and **Supplementary Table 4** for details and an extensive list of references on these genes). Briefly, we divide the NMB5000 genes into five groups:

Group 1 (n = 12) contains genes that have strong statistical genetic evidence linking them to human cancers through point mutations and short insertions and deletions. Seven genes (*ERBB3, EZH2, PIK3CA, PIK3R1, RASA1, SOS1, STK11*) are in the Cancer5000 set previously published in Ref. 1 (**Supplementary Table 4**). The convergence on seven genes between these two independent methods (which is highly significant at P = 1.9e-5) serves as an overall validation of the NMB approach (see **Methods**). The five remaining genes (*ESR1, AFF2, AKT3, PIK3R2, PIK3CB*) serve as an additional validation of the NMB predictions because they can also be considered a set of true positive cancer genes independent of the Cancer5000 set. One example is *ESR1* which is significantly mutated in hormone sensitive breast cancer. Another is *AFF2*, which plays an unknown role in cancers. Our analysis shows that the corresponding protein is in a high-confidence network with PTEN, BRAF, KRAS, CTNNB1 and GRB2, which at the gene level have pan-cancer mutations at varying levels of significance (**Fig. 3a**) which leads to an NMB q = 0.07 for this gene. Interestingly, a recent study has identified this gene as significantly mutated in breast tumours^16^, and in the same publication other members of the AFF2 network are linked to breast cancer. For example, *PTEN* is also significantly mutated and *KRAS* and *BRAF* are amplified in 32% and 30%, respectively, of the patients in this cohort through copy number variants. Another interesting gene is *PIK3CB* which has a NMB q = 0.016 (**Fig. 3b**). *PIK3CB* has been suggested to play an important role in PTEN-deficient tumours^17,18^ and it has been shown that both in cell lines and in vivo down-regulation of *PIK3CB* leads to the inhibition of tumour growth^19^. Furthermore, overexpression of *PIK3CB* can transform cells in vitro^20^. In our analysis the corresponding protein PIK3CB is in a network with KRAS, ERBB2, EGFR, MTOR, PIK3R1, RAC1, AKT1 and HRAS. The function of PIK3CB is suggested to be related to DNA synthesis/replication and cell mitosis^21^ and there is further evidence that mutational activation can enhance basal association with membranes and thereby increase proliferation and survival^22^. In a recent study of advanced prostate cancer, *PIK3CB* was found to harbour (likely activating) mutations as also be part of fusion-events leading to increase expression^23^. In functional validation experiments that we describe in a distinct manuscript (E. Kim, et al., in preparation), it has been shown that over-expression of mutant alleles of PIK3CB (E497D and A1048V) identified in the pan-cancer sequencing study^1^ drives subcutaneous tumour formation in mice further suggesting its likely causal link to human cancer.

Group 2 (n = 8) contains genes with genetic evidence linking them to cancer risk in patients, but which have not yet been identified to have point mutations in large cancer sequencing studies (e.g., *CBFA2T2*, in which the t(8;21)(q^22^;q^22^) translocation is one of the most common karyotypic abnormalities in patients with acute myleoid leukemia). Another gene in this group is *E2F4* (**Fig. 3c**, NMB q = 0.03), which encodes a transcription factor that plays a critical role in cell cycle regulation and binds to three known tumour suppressors, pRB, p107 and p13. When affected by microsatellite polymorphisms, *E2F4* is linked to an enhanced risk for developing breast cancer^24^. Interestingly, it has also been shown that loss of *E2F4* can suppress tumourigenesis^25^ while its expression is needed for the proliferation of e.g., colorectal tumours^26^. In our work we show that it is in a network with e.g., MGA, which is associated with breast cancer^1^.

Interestingly, it has been shown in heterozygous Rb loss of function mouse models that E2F4 loss suppresses the development of both pituitary and thyroid tumours suggesting a direct regulatory relationship between these proteins. Finally, the E2F4 network contains SMAD4, which is found to be mutated in**∽** 10-35% of all colorectal cancers^27,28^.

Group 3 (n = 15) contains genes linked with strong functional evidence to cancer processes and networks in animal or cell models, but where human genetic data (mutations, copy number variants or karyotypic abnormalities) does not yet strongly incriminate these genes in any patient. For example, *ETV7* is functionally linked to leukemia and accelerates lymphoma development in transgenic mice^29^. Another candidate, *PSEN1*, is linked to the down-regulation of WNT. Loss-of-function mutations in *PSEN1* might lead to the over-activation of WNT and promote tumourigenesis in colon cancer^30,31^.

Group 4 (n = 22) contains genes that have been linked to cancer through e.g. aberrant expression, but where the direct functional evidence is not as conclusive as in group 3 (e.g., BMX which is upregulated in human prostate cancer). For example, RUNX2 (**Fig. 3d**, NMB q = 0.07) is a transcription factor modulating e.g. the activities of both RNA polymerases I and II^32^.

**Figure 2.**
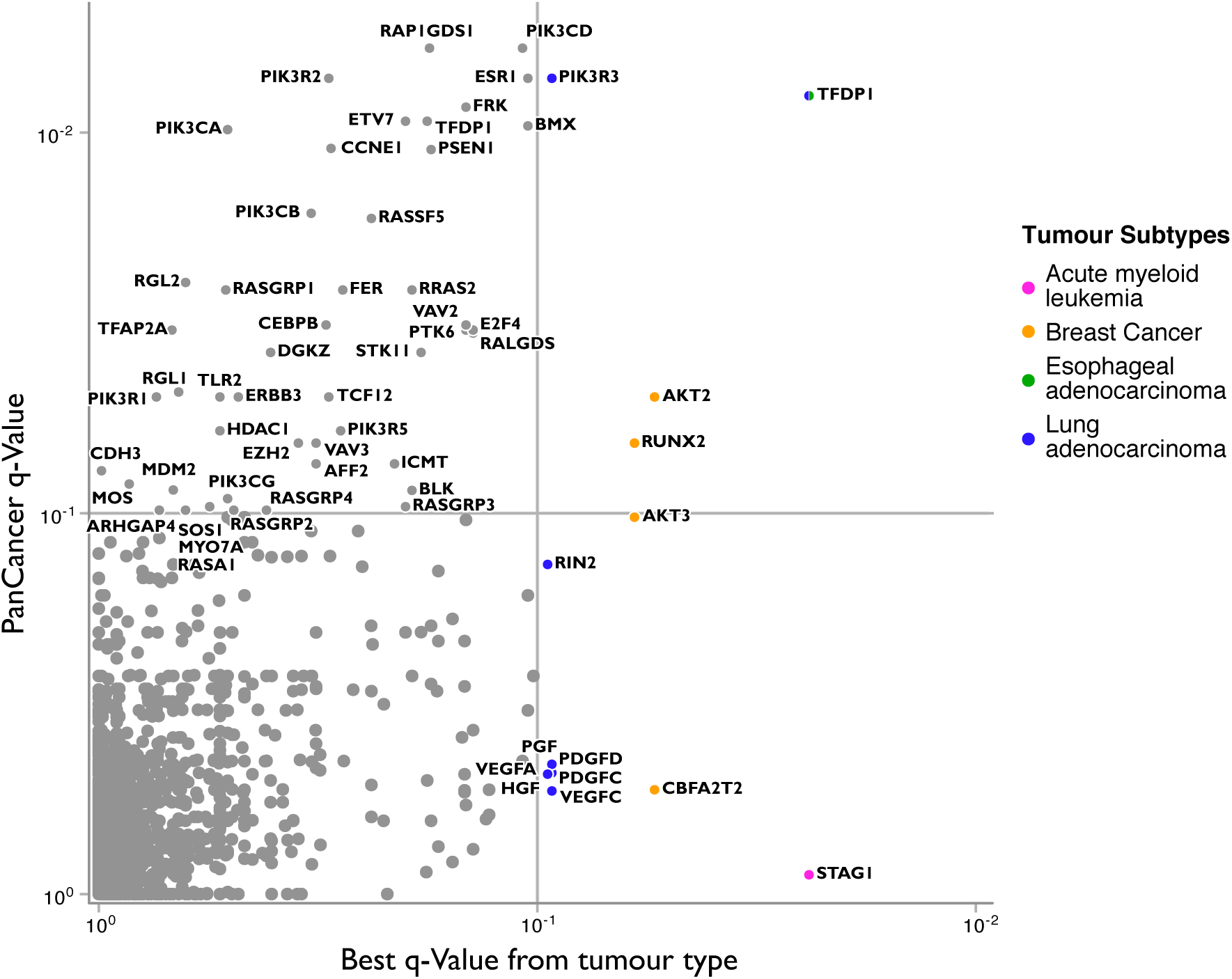
Sixty-two NMB-imputed cancer genes. Genes are represented as individual dots and plotted along the x-axis by the NMB q value from the most significant of 21 tumour types, and on the y-axis by the NMB q value when 4,724 tumours are analyzed as a combined pan-cancer cohort. Significance at FDR q *<*= 0.1 is indicated on each axis by grey lines. Genes above the horizontal line are significant in the pan-cancer analysis. Genes to the right of the vertical line are significant in at least one tumour type with the most significant tumour type indicated by the node color. Genes in the upper right quadrant are significant in both the pan-cancer data and in an individual tumour type.

**Figure 3.**
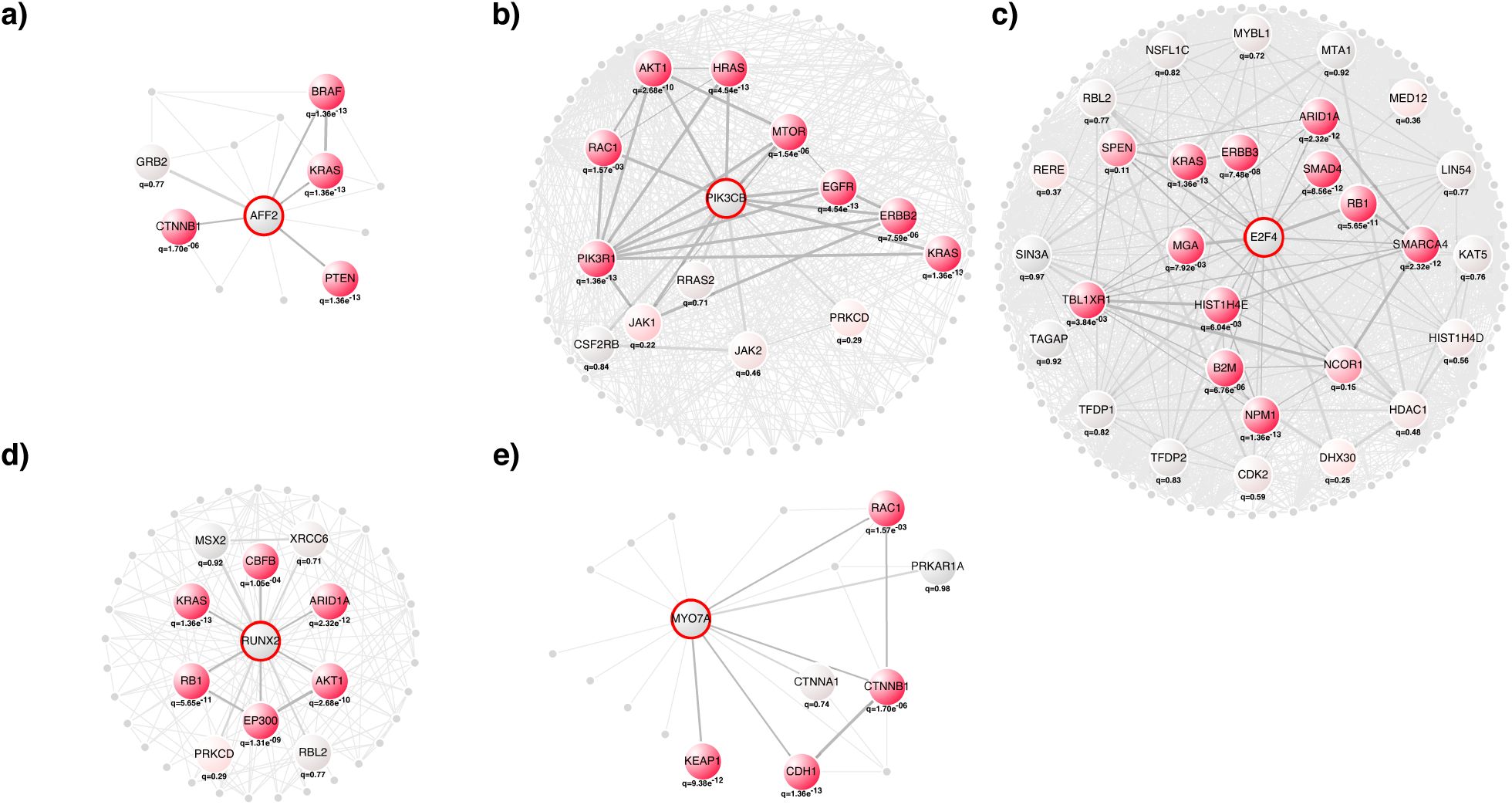
Examples of NMB-imputed cancer genes. The first order network of **a)** AFF2 (dark grey with red border, NMB q = 0.07), **b)** PIK3CB (dark grey with red border, NMB q = 0.016), **c)** E2F4 (dark grey with red border, NMB q = 0.03), **d)** RUNX2 (dark grey with red border, NMB q = 0.07), and **e)** MYO7A (dark grey with red border, NMB q = 0.06) that are significant in the NMB analysis. Large nodes other than AFF2, E2F4, PIK3CB, RUNX2, and MYO7A are colored by the significance of the pan-cancer q value of the corresponding gene, where light grey or no shading represents q close to 1 and red q *<<* 1, with darker red representing more significant q values as indicated. Small nodes represent genes with q = 1 or not annotated in the pan-cancer data. These examples illustrate the diverse functional and genetic architectures of cancer networks that can lead to significant NMBs and highlight the importance of considering, and normalizing for, these architectures in the process of calculating statistical significance (**Supplementary Note 1**). Interactive networks with names and q values for all proteins can be seen on http://www.lagelab.org.

It is significant in the NMB pan-cancer analysis, but also in the tumour-specific NMB analysis using only breast cancer data. As for all genes in this group there is no direct genetic evidence for its role in human cancer, but it has been shown that induction of *RUNX2* enhances tumour invasiveness in prostate cancer^33^, while silencing through siRNA prevents cell invasion for thyroid cancer^34^ and colorectal cancer^35^ cells. As expected, many proteins in its neighbourhood (CBFB, KRAS, RB1, ARID1A and AKT1) are associated with breast cancer^1^. Moreover, genes in the neighbourhood of RUNX2 are significantly mutated in many tumour types (seven, six, and six, for ARID1A, KRAS and RB1, respectively), suggesting this is an important network across many tumour types.

Group 5 (n = 5) includes genes for which we could not find any connection to cancer in the literature. One example is MYO7A (**Fig. 3e**), which encodes an unconventional myosin, that is plays a role in cell adhesion^36^ and is most famous for its role in Usher syndrome^37^. Nonetheless nonmuscular myosins are known to be involved in tumour progression, cancer cell invasion, and metastasis^38^, and in our work MYO7A is shown to physically interact with e.g., the cadherin/catenins complex (**Fig. 3e**) leading to a NMB q = 0.06. Specifically, MY07A interacts with *β* -Catenin (CTNNB1) and cadherin 1 (CDH1) in InWeb (source data from^39^). Where somatic *β* -Catenin mutations are known to be involved in colon and other cancers^40^, germline mutations in cadherin 1 are known to be a major cause of familial gastric cancers^41^. Further interactions of MY07A with RAC1 (a known Rho GTPase that regulates the functions of the cadherin complex) strengthen this link and suggests an interesting potential role for MY07A in tumourigenic processes linked to the cadherin function and WNT signaling.

### The genetic architecture of cancer networks across 21 tumour types

The reason we only identify significant NMB-imputed genes in 4 out of 21 (*<*20%) of tumour types could either be that the genetic data in the remaining 17 tumour types is not powered to lead to NMB significances at a FDR *<*0.1 or it could indicate a fundamental difference between cancer types in terms of functional and genetic architecture (i.e, that genes driving some cancers form highly mutated networks that can be captured using functional protein networks while others do not). To test these two hypotheses we iterated over the 21 tumours and first calculated tumour-specific NMB scores. Second, we defined a set of driver genes known to be significant in that tumour type and for tumours with more than four driver genes measured the performance of the tumour-specific NMB in terms of distinguishing that set of driver genes from all other genes represented in the InWeb network. Third, we compared the performance of the tumour-specific NMB to the performance of NMB scores derived from the 20 other tumours and from the pan-cancer data (**Fig. 4**).

For example, we assembled a set of driver genes from breast tumours (BRCA) by identifying genes significantly mutated in this tumour type in Ref. 1. We used mutation data from this tumour type to derive NMB_BRCA_ scores and measured their classification performance on the BRCA driver genes, which they could accurately distinguish with an AUC = 0.76. This result was compared to the ability of NMB scores derived from the 20 other tumour types as well as the a pan-cancer dataset calculated after excluding data from the BRCA cohort (NMB_pan-cancer minus BRCA_) to classify the BRCA driver genes (**Methods**). The NMB_pan-cancer minus BRCA_ scores increased the ability to accurately classify BRCA driver genes slightly to an AUC of 0.77, while the NMB scores derived from the 20 other tumour types all performed worse on BRCA driver genes than the NMB_BRCA_ (**Fig. 4**, ranging from an AUC of 0.75 [bladder] to an AUC of 0.47 [prostate], and median AUC = 0.69).

In 9 of 17 (or 53%) of tumour types we see that the tumour-specific NMB approach can accurately classify genes significantly mutated in that tumour type with an AUC *>*0.7 (**Fig. 4**). In 10 of 17 of tumours the NMB scores based on pan-cancer data (calculated excluding data from the tumour in question) outperforms the NMB scores based on the specific tumour being tested (on average the predictive power of the pan-cancer NMB is 8% better than the tumour-specific NMB), and it is a consistent observation that the tumour-specific NMB score is better at classifying driver genes from the corresponding cancer than NMB scores derived from any of the 20 other tumours as expected. Remarkably, close to 60% (or 10 out of 17) of the tumour types the driver genes can be classified with an AUC *>*0.70 either by the tumour-specific NMB or the NMB_pan-cancer minus X_ that excludes data from tumour X being analyzed.

These observations highlight that, despite our inability to measure significant genes in the 80% of tumours using the NMB approach, there is clear evidence that driver genes across most types of cancer sequenced to date assemble into networks that have a non-random mutation burden and suggests a way to quantify the pathway convergence across distinct tumour types. Our results also stress the value of meta-analyzing multiple tumour types into pan-cancer data sets to increase the statistical power to find convergent pathways and networks that are of wide interest to unraveling the biology across cancers.

### NMB significances are independent of mutation frequencies

Current statistical methods for identifying significant genes based on cancer-sequencing data (e.g., the MutSig suite) have more power to detect genes with high frequency mutations in cancer patients versus those with intermediate and low frequencies^1^.

We hypothesized that significances assigned to index genes based on the NMB method are not dependent on its mutation frequency, because the signal is tallied across the gene’s first order network and therefore is independent of mutation frequencies in the index gene itself. In fact, the index gene may not be mutated at all in any tumour sequenced to date. We plotted the distribution of mutation frequencies as the function of P-values determined from the pan-cancer data by the MutSig suite and NMB (**Fig. 5a**), and confirmed that former are correlated with mutation frequencies while the latter are not.

**Figure 4.**
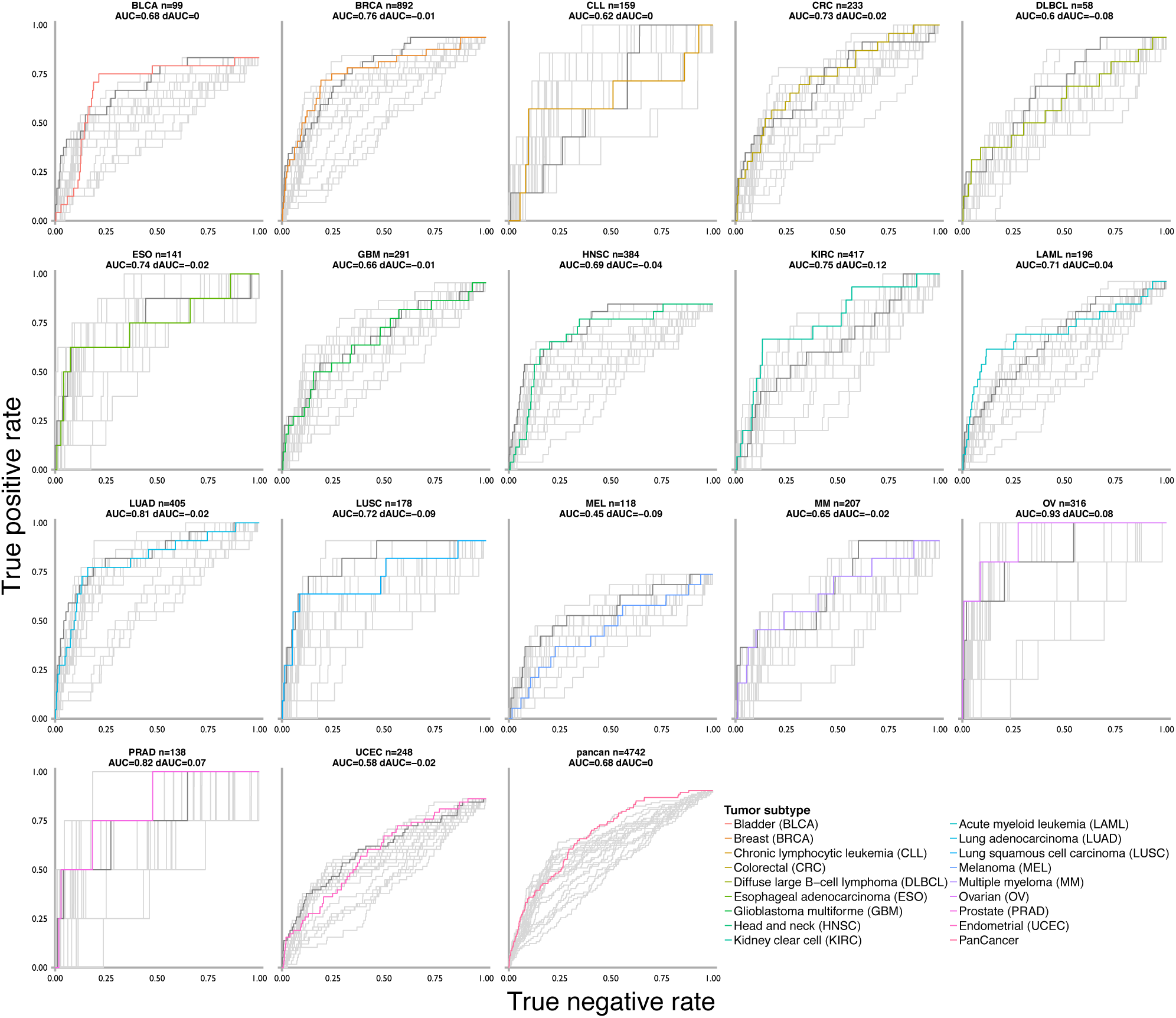
A quantitative analysis of cancer networks across 21 tumour types. In this figure, each box illustrates the results from the tumour-specific NMB analyses of 17 tumour types that have at least four defined driver genes (first 17 boxes) or for pan-cancer genes (last box). In each plot, for example for breast cancer [BRCA, second box], the AUC calculated with the NMB score corresponding to mutation burdens from BRCA (NMB_BRCA_) is indicated by the orange line. The ability to distinguish BRCA driver genes using an NMB score derived from pan-cancer data calculated after excluding the BRCA data (NMB_pan-cancer minus BRCA_) is indicated with a dark grey curve. The lighter grey curves indicate the ability of NMB scores derived from the 20 other tumour types to accurately classify BRCA driver genes. The difference in performance using NMB_pan-cancer minus BRCA_ and NMB_BRCA_ (in this case 0.76-0.77 = -0.01) is indicated above the plot as the differential AUC (or dAUC). Although the pan-cancer data is better at classifying BRCA driver genes than the BRCA-specific mutation data, the NMB_BLCA_ is better at classifying BLCA driver genes than NMB scores derived from any of the 20 other tumor types (light grey curves, median AUC = 0.69).

To further explore this phenomenon, we plotted the relative proportion of high frequency, intermediate frequency, and low frequency genes amongst the candidate cancer genes predicted by the MutSig suite and NMB both for the pan-cancer analysis, and for the Cancer5000 and NMB5000 sets. Using the pan-cancer data, the relative proportion of low frequency candidate cancer genes identified with the NMB approach versus the MutSig suite is more than three times higher (13% vs 4%). This enrichment is mirrored in the NMB5000 vs. Cancer5000 sets, where the proportions of low frequency genes are 15% vs. 4%, respectively (**Fig. 5b**), suggesting that approaches like the NMB can complement existing methods and contribute independent signal to finding potentially important cancer genes with low-frequency mutations in the tail of mutation distributions from tumour sequencing studies.

**Figure 5.**
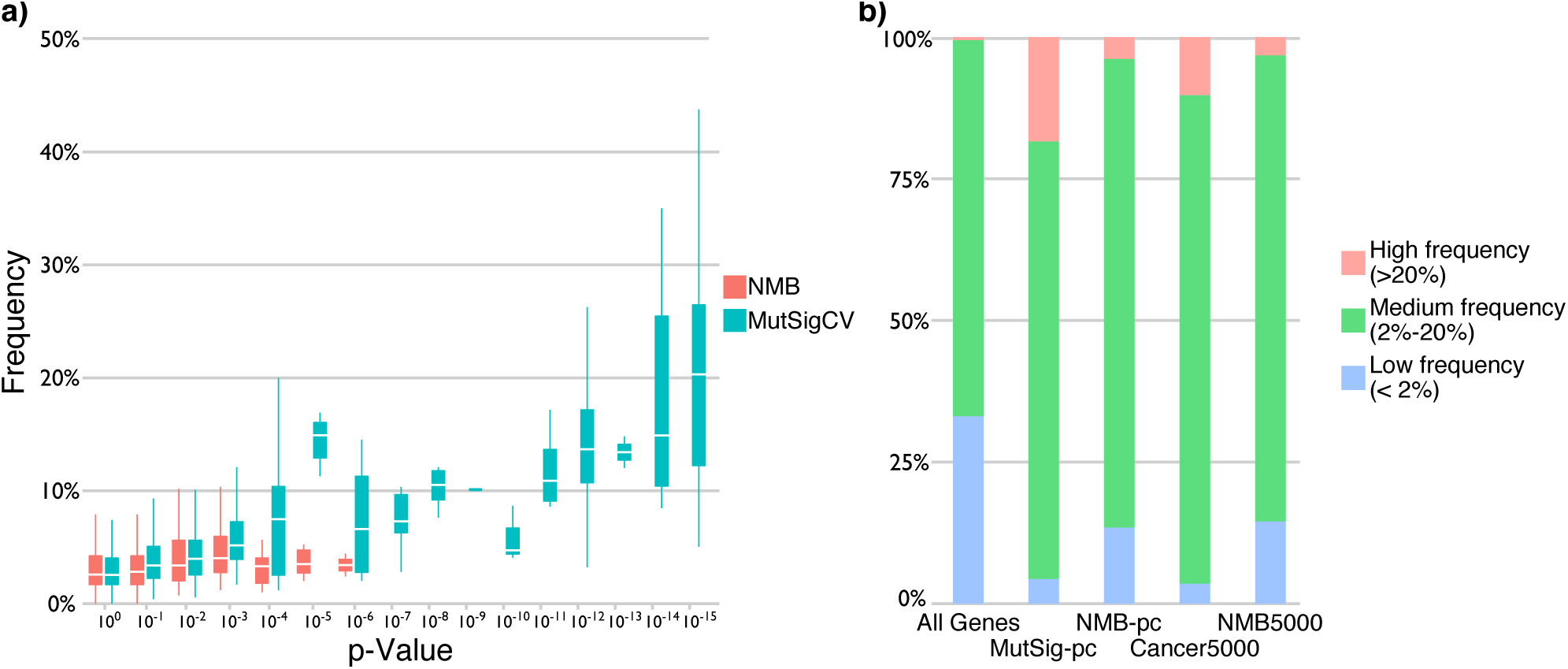
Network mutation burdens significances are independent of mutation frequencies. **a)** A box plot of the NMB (red) and MutSig suite (blue) P values (x-axis) versus mutation frequency distributions (y-axis). Boxes represent median first and third quartile of the frequency distribution for a given P-value bin (NMB values are permutation-based which limits us to deriving P *>*= e^-^6). In contrast to the MutSig suite, NMB P values are not correlated with mutation frequencies. **b)** The mutation frequency of all (18,600) genes in the genome is shown in the left column colored as indicated in the legend. Columns two and three show that relative to the total amount of genes classified as significant in the pan-cancer data, the NMB analysis (NMB-pc) identifies 3 times more genes with low frequency mutations than the MutSig suite (MutSig-pc). These proportions are slightly higher when comparing the NMB5000 and Cancer5000 data sets (columns four and five).

### Networks lead to molecular clues in patients without established driver mutations

In 17 of the 21 cancer studies recently described in Ref 1, there is at least one patient that does not have mutations in the genes established to be drivers in the cancer and publication in question. To test if the NMB approach can contribute to providing molecular clues in these patients, we downloaded the mutation profiles (**Supplementary Table 5**), and compared the proportions of patient-derived mutations classified as damaging by PolyPhen2^42^ in the NMB5000 gene set, to a random expectation (i.e., the same proportions in all genes in the genome, **Fig. 6a** and **b**). We see a significant enrichment of damaging mutations in the NMB5000 genes (P = 0.016 using Fisher’s exact test to make a categorical comparison and P = 0.046 using a non-parametric two sample Kolmogorov-Smirnov test to compare distributions of continuous mutation scores). Moreover, the proportions of damaging or deleterious mutations across NMB genes in patients without known driver mutations are remarkably similar to the analogous proportions in the Cancer5000 set (**Fig. 6c** and **d**), but, as expected, the results are many orders of magnitude more significant due to many more genes in the Cancer5000 set.

The four NMB-imputed cancer genes with the highest ratio of damaging to benign mutations in the patients are *PIK3CA, MY07A, PIK3R5*, and *PDGFD* (see additional details on specific mutations, cancers, and these genes in **Supplementary Table 6**). *PIK3CA* is a well-known oncogene in breast cancer^43^, which showed damaging mutations in patients with colorectal cancer.

*MY07A* (**Fig. 3d**) is a gene not previously linked to cancer and has damaging mutations in patients with colorectal cancer, esophageal adenocarcinoma, kidney clear cell carcinoma, lung squamous cell carcinoma, melanoma, and neuroblastoma. *PIK3R5* is a subunit in the Phosphoinositide-3-Kinase pathway linked to cancer through several other members (Lawrence et al.1) which has damaging mutations in patients with kidney clear cell carcinoma, neuroblastoma and prostate cancer. Lastly, *PDGFD* showed damaging mutations in patients with colorectal and kidney clear cell carcinoma.

Interestingly, of the five NMB-imputed cancer genes we point to in **Fig. 3**, three are amongst the ten genes with the highest ratio of damaging to benign mutations in patients (**Supplementary Table 6**). In fact, *MY07A* (**Fig. 3e**) has the next-highest proportion (7 fold more damaging mutations than benign mutations) after *PIK3CA* (with 8.5 fold more damaging than benign mutations). The ratio for *MY07A* is significantly higher than we would expect by random (nominal P = 0.0092 and adjusted P = 0.046 after correction for testing all five genes from **Fig. 3**, see **Supplementary Figure 6** for details). These observations make it an interesting candidate to follow up in particularly colorectal and esophageal cancers.

We further compared the ratio of damaging to benign mutations in the 55 NMB5000 genes that do not overlap with the Cancer5000 set to the distribution of the same proportions in 100 sets of 55 randomly chosen Cancer5000 genes (i.e., matched and downsampled sets). Here, the proportion of damaging to benign mutations is comparable to the expectation from Cancer5000 genes (i.e., within the 95% confidence interval of the mean, **Supplementary Figure 7**). We repeated this analysis for each of the five literature curation groups individually showing that the proportions for Groups 1 and 5 are comparable to matched and downsampled sets of Cancer5000 genes (within the 95% confidence interval of the mean, **Supplementary Figure 8**). Together this provides support for the cancer relevance of the 55 cancer genes we identify exclusively through the NMB approach as well as groups 1 and 5 even when they are considered as individual sets.

**Figure 6.**
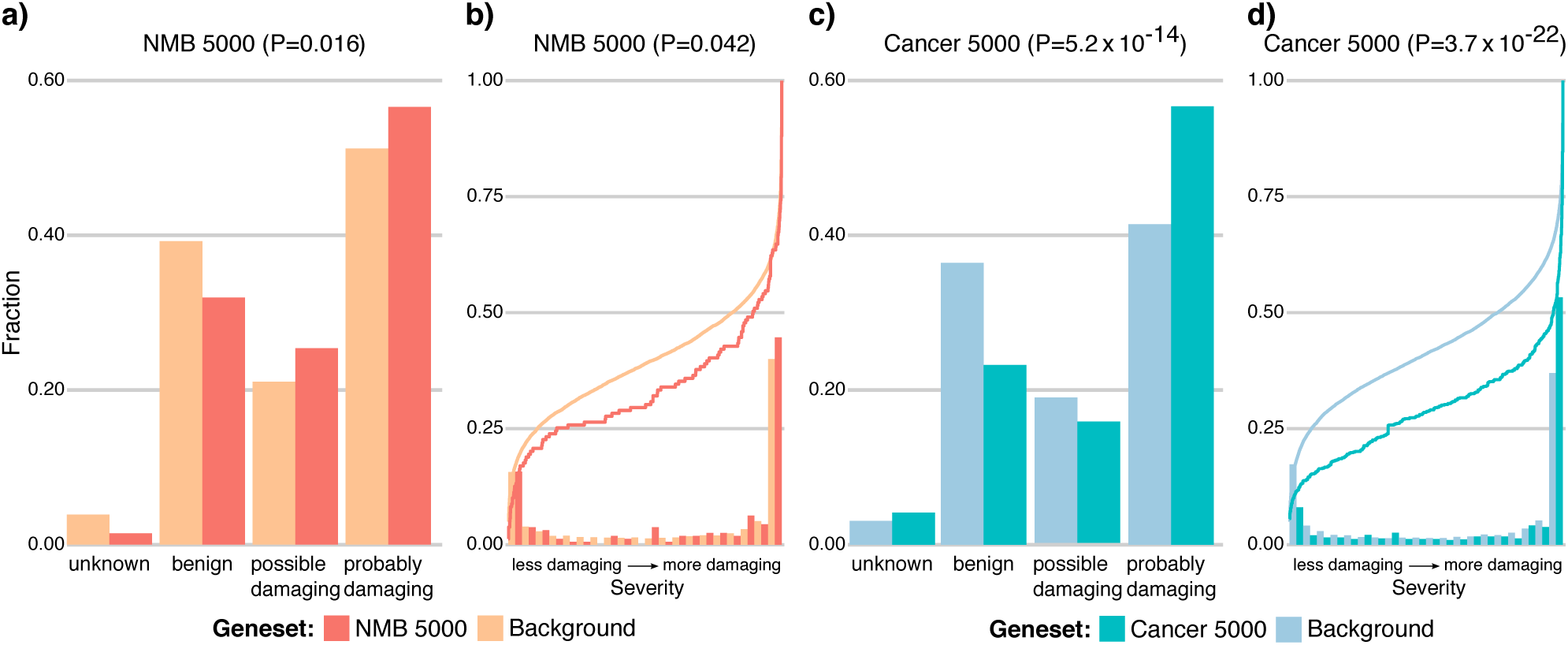
Cancer patients without established driver mutations are enriched for deleterious mutations in NMB5000 genes. **a)** We compared the fraction of genes with damaging (i.e, probably damaging and possibly damaging pooled into one set) versus benign mutations (as determined by PolyPhen2) in the NMB5000 genes on the background of all genes in the genome, and show a statistically significant enrichment of damaging mutations in the NMB5000 set (P = 0.016, using Fischer’s exact test, NMB5000 is indicated by dark red and all genes in the genome by light red). **b)** Using PolyPhen2, we transformed all mutations observed in the NMB5000 set to continuous normalized scores of how much the mutation is predicted to affect gene function negatively (less damaging to more damaging oriented left to right on the x-axis). When comparing to all genes in the genome, mutations in the NMB5000 genes are significantly depleted for less damaging PolyPhen scores, and significantly enriched for more damaging PolyPhen scores (P = 0.046, using a non-parametric two-sample Kolmogorov-Smirnov test, and histograms show the binned proportions, the line the cumulative distributions of scores). For comparison, we show the results for the same analyses on the Cancer5000 set in panels **c)** and **d)**, respectively with Cancer 5000 significant genes in dark blue and the background genes in light blue. While the trends and proportions of deleterious versus benign mutations observed in the Cancer5000 genes are similar to our observations for the NMB5000 genes, thus supporting the cancer relevance of the NMB5000 set, the statistical significances levels are higher due to more genes in the Cancer5000 set.

## Discussion

Here we quantify and compare the degree to which cancer genes across 21 tumour types can be accurately classified based only on the pan-cancer mutation burden in their first order functional protein network (which excludes the gene itself). Data on somatic mutations was acquired from the exome sequences of 4,742 human cancers and their matched normal-tissue samples across 21 tumour types1. We found that very well established cancer genes can be accurately classified based on their NMB score and we show that it is a general principle across most cancers that driver genes form networks that can be explored using proteinprotein interaction data. Our results indicate that network-based approaches such as the NMB are a scalable and cost-efficient way to extract more information from existing cancer geneomes in a purely compuatational manner and that they can contribute to a deeper understanding of tumour biology widely across indications.

To test the general applicability of our NMB approach and to investigate if candidate cancer genes could be robustly predicted in a range of different functional genomics networks using the statistical framework we have developed, we repeated our analysis in gene networks based on mRNA coexpression, gene coevolution profiles, cancer synthetic lethality relationships, and cell perturbation profiles. While we observe the strongest signal in the protein-protein interaction network data from InWeb (**Fig. 1**), there is evidence that significant cancer networks can be detected in three of four networks using the NMB approach (**Supplementary Figure 6**).

It is important to stress that our analysis does not guarantee that a gene with a significant network mutation burden is a cancer gene which is why we coined the term NMB-imputed cancer gene. However, we find conclusive evidence that cancer genes across 10 of 17 tumour types have a non-random mutation burden in their immediate functional protein interaction neighborhood and that this signal is strong and consistent enough to enable an accurate classification of driver genes involved in many of these cancers. We illustrate that this phenomenon is not significantly driven by “knowledge contamination”, meaning that our approach performs comparably even when the most established cancer genes (and those with the highest mutation frequencies) are disregarded in the calculation of NMB. Importantly, we observe a comparable signal for very well established cancer genes and cancer genes emerging from the newest unbiased cancer sequencing studies that we know little about in terms of biology. While it is only possible to classify the NMB-imputed cancer genes we point to here as bona fide functional cancer genes based on rigorous experimental follow-up analyses, we make a comprehensive comparison and quantification of the signal coming from the gene network architecuture in cancers. We furthermore illustrate how this principle can be leveraged into a robust statistical framework that complements gene-based tests to generate new hypotheses about possible driver genes from existing sequencing studies. Our approach should become more and more powerful as sequencing studies increase in size in the future and as functional genomics networks increase improve in coverage and accuracy.

We predict 62 genes to have a significant mutation burden in their functional molecular neighborhood. We call this set of genes the NMB5000 set and show it significantly converges of the same genes as the Cancer5000 set from the recent MutSig suite analysis1. One gene identified by our approach is *PIK3CB* and in functional validation experiments (described E. Kim et al., manuscript in preparation) we show that over-expression of mutant alleles of *PIK3CB* (E47D and A1048V) identified in the pan-cancer sequencing study (Ref. 1) drive subcutaneous tumour formation in mice. This example demonstrates that combining NMB predictions with follow-up experimental data can detect novel cancer genes. Furthermore, our analysis suggests *MY07A* as an interesting candidate cancer gene that potentially can play a role in colorectal and esophageal cancers. The NMB5000 set also contains genes known to play a role in cancers through copy number changes (*CCNE1, TFDP1* in breast cancer), by fusing with other genes (*RAP1GDS1* in T-cell acute lymphoblastic leukemia^44^, *CBFA2T2* also known as *MTGR1* in Acute myeloid leukemia^45^) or by being affected by microsatellite polymorphisms (*E2F4*).

We further show that the significances assigned to individual genes by the NMB approach are independent of mutation frequencies, a limitation of existing sequenced-based methods, and we observe an enrichment of damaging mutations in clinical samples for genes in the NMB5000 set further supporting that genes identified using this method are relevant to cancers and point to new tumour-relevant biology. Simultaneously, our analysis revealed 27 genes (or 44% of all NMB-imputed cancer genes) that were not previously strongly linked to cancers (i.e., placed in groups 4 and 5 of our literature review). These genes may provide new clues towards a comprehensive understanding of tumour biology. Towards this aim all network data are deposited on http://www.lagelab.org as a resource for the community along with graphical displays of the individual networks, null distributions of composite mutation burden for the network in question, and mutation information on genes in the networks that drive the NMB signal.

The methodological framework developed here is flexible and could easily be extended to any user-defined functional genomics network, but also to include many different types of mutation data (e.g., structural variation) or to be integrated as an independent signal with the MutSig suite of tools to increase power to detect cancer candidate genes. We expect that with better interaction networks and larger collections of sequenced tumours, approaches leveraging the principles we describe and quantify here will become increasingly powerful and applicable for biological discovery in cancers.

## Methods

### Calculating the network mutation burden

For a given index gene the network mutation burden (NMB) is formalized into a probabilistic score that reflects the index-gene-specific composite mutation burden [i.e. the aggregate of single-gene MutSig suite q values from Lawrence et al.^1^] across its first order biological network and is calculated via a three-step process: First, we identify all genes it interacts with directly at the level of proteins, only including high-confidence quality-controlled data from the functional human network InWeb^14,46^, where the vast majority of connections stem from direct physical interaction experiments at the level of proteins. Second, the composite mutation burden across members of the resulting network is quantified by aggregating single-gene MutSig suite q values from Lawrence et al.1 into one value *ϕ* using an approach inspired by Fisher’s method for combining p-values:

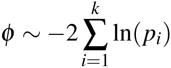

Where *p*_*i*_ is the MutSig suite q value for gene i, and k is the amount of genes in the first order network of the index gene (i.e. the index gene’s degree). Third, by permuting the InWeb network using a node permutation scheme, we compare the aggregated burden of mutations *ϕ* to a random expectation. In this step, the degree of the index gene, as well as the degrees of all genes in the index gene’s network is taken into careful consideration. The final NMB score of an index genes is therefore an empirical P value that reflects the probability of observing a particular composite mutation burden across its first order physical interaction partners (at the level of proteins) normalized for the degree of the index gene as well as the degrees of all of its first order interaction partners. Because we are interested in estimating the mutation burden independent of the index gene, this gene is not included in the analysis and it does not affect the NMB calculation meaning that for any given gene MutSig suite significances are independent of NMB significances (i.e., the Cancer5000 gene set and the NMB5000 gene set are independently predicted). We tried several ways of calculating a networks composite mutation burdens (step 2) and permutation methods (step 3) which all give similar results (**Supplementary Note 1** and **Supplementary Figure 2**) illustrating the robustness of our approach.

**Classifying cancer genes.** For each gene represented in InWeb (12,507 or 67% of the estimated genes in the genome), we used the gene-specific NMB probability to classify it as a cancer candidate gene or not. True positive genes were defined as the Tier 1-4 genes (described in the main text and **Supplementary Table 1**) pan-cancer genes from Lawrence et al1. Tier 5 (**Supplementary Table 1**) was included as a negative control. True negatives were defined as all genes in InWeb that were not in Tiers 1-5 which is likely conservative as many of these might be yet undetected cancer genes. We used the NMB probability as the classifier and calculated the AUC for each Tier. We used both a node permutation scheme and a network permutation scheme (discussed in detail in **Supplementary Note 1**) to generate random networks and repeated the analysis to assess the significance of the reported AUCs. The results were not significantly influenced by choice of permutation method supporting the robustness of the NMB concept and our chosen approach (**Supplementary Note 1** and **Supplementary Figure 2**).

### Predicting new NMB-imputed candidate cancer genes

To predict new NMB-imputed cancer genes we used a node permutation scheme to create 106 permuted networks. NMB probabilities were determined for every gene in InWeb that was covered by interaction data. The FDR q values were calculated as described by Benjamini and Hochberg based on the nominal P values controlled for 12,507 hypotheses. We performed NMB analyses with the pan-cancer q values, as well as q values from each of the 21 tumour types for which they were available. As it is a technical limitation of the NMB approach that it is currently not possible to make 5.5 * 10^6^ network permutations we could not create a dataset where we correct for all 12,500x22 hypotheses tested in the NMB5000 set. For that reason our work does not have the equivalent of the Cancer5000-S (the stringent) set from Lawrence et al.^1^ where the control for all of the hypotheses is carried out simultaneously.

### Enrichment of deleterious mutations in patients without established driver mutations

Genome variation data for patients was manually extracted from the variant calling tables of TCGA sequencing studies (**Supplementary Table 1**). If a patient group without known driver mutations was already defined by the authors of the study, patients were extracted directly from the variant calling file (this was the case for rhabdoid tumour, medulloblastoma and neuroblastoma). For studies with no predefined patient groups of this type, we identified patients similarly to the four articles above as those with no damaging mutations (frame shift or missense) in any of the genes significantly mutated in the article in question. Variants from the studies were either aligned using NCBI Build 36 or 37 of the human genome depending. To translate coordinates reported for build 36 to build 37, the tool liftOver from the UCSC Genome Bioinformatics Site (http://hgwdev-kent.cse.ucsc.edu/) was used. The ENSEMBL Variant Effect Predictor (http://www.ensembl.org/info/docs/tools/vep/index.html) was used to analyze all reported variants. Briefly, we extracted the precompiled PolyPhen242 (http://genetics.bwh.harvard.edu/pph2/) predictions. For genes with multiple transcripts, we selected the most damaging transcript effect for the analysis and P values were calculated using a non-parametric two-sample Kolmogorov-Smirnov and the Fisher’s exact test for distributions and mutation categories, respectively. The background distribution was defined as all genes represented in InWeb, but not significant in the NMB analysis.

